# Frequency and Distribution of Corneal Astigmatism and Keratometry Features: Methodology and Findings of the UK Biobank Study

**DOI:** 10.1101/654236

**Authors:** Nikolas Pontikos, Sharon Chua, Paul J Foster, Stephen J Tuft, Alexander C Day, UK Biobank Eye and Vision Consortium

## Abstract

**Purpose:** To describe corneal astigmatism in the UK Biobank population, to look for associations with other biometric variables and socio-demographic factors, and to report the proportion with abnormal keratometry and irregular astigmatism suggestive of pathological corneal ectasias such as keratoconus.

**Methods:** Cross-sectional data were obtained from UK Biobank (www.ukbiobank.ac.uk/). A subsample of 107,452 participants from UK communities had undergone an enhanced ophthalmic examination including autorefractor keratometry (Tomey RC 5000, Tomey Corp., Nagoya, Japan). After quality control and applying relevant exclusions, data on corneal astigmatism on 83,751 participants was available for analysis. Potential associations were tested through univariable regression and significant parameters carried forward for multivariable analysis.

**Results:** In a univariable analysis, the characteristics significantly protective against corneal astigmatism were gender (male), older age, darker skin colour and increased alcohol intake (all p<0.001). The parameters significantly associated with increased corneal astigmatism were older age at completion of full time education, use of UV protection and lower corrected visual acuity. After inclusion in the multivariable analysis, age, gender, age at completion of full time education, corrected visual acuity and skin colour remained significant (all p<0.001). Increased corneal astigmatism was also found to be significantly associated with amblyopia or strabismus. No individuals with abnormal keratometry or irregular astigmatism were reported.

**Conclusions:** This analysis of associations with astigmatism in a large cohort of volunteers confirms previous associations including adverse associations with younger age and female gender, and identified novel associations including darker skin colour and frequency of alcohol intake. The highest risk group for corneal astigmatism were younger females of lighter skin colour, having completed full time education later, with higher logMAR corrected visual acuity. We also confirmed that corneal astigmatism is a high risk factor for amblyopia and strabismus. Finally since no cases of keratoconus were identified, this would suggest that simple keratometry indices may not be sufficient for population screening of keratoconus.

## Introduction

Uncorrected refractive error is the leading cause of moderate to severe visual impairment in all age groups globally (101.2 million individuals in 2010), and the second commonest cause of avoidable blindness in children after cataract (6.8 million individuals in 2010) [1][2]. Refractive error (ametropia), is a significant public health burden, frequently associated with worse visual acuity and higher risk of developing amblyopia (lazy-eye). Corneal astigmatism is a leading cause of refractive error in children and a significant increase in myopia as well as astigmatism has been reported in the Singaporean population[3] over the last 12 years. In a previous study, Cumberland et al [4] found that 54% of participants in the UK Biobank (UKBB) had a refractive error.

The two major components of refractive error in the eye are astigmatism, which is linked to the refractive properties of the cornea and the lens, and the spherical refractive error, which is additionally related to axial length of the eye. Spherical refractive error (myopia or hyperopia) can be corrected by a spherical spectacle lens. Astigmatism is caused by a corneal component and lenticular component and can in many cases be corrected by a cylindrical spectacle lens. Corneal astigmatism occurs when there are differences in the radius of curvature of the cornea in different meridians such that there is a different focal point for each meridian, with an area of intermediate focus between the two termed the conoid of Sturm[5]. In regular astigmatism the strong and weak meridians are normally at 90 degrees to each other. Regular astigmatism is usually defined according to the orientation (meridian) of the steepest radius of curvature of the cornea in relation to the horizontal axis of the cornea (0 to 180 degrees). The axis of regular astigmatism is normally at 90 degrees to the steep meridian, and this angle also corresponds to the minimum radius of curvature of the cornea. Its axis defines the orientation of the negative power cylindrical spectacle lens required to cancel the effect of the astigmatism. Irregular astigmatism occurs when the radius of curve of the central cornea in any one meridian varies, and the angle between the strong and weak meridians may not be 90 degrees. In irregular astigmatism, full correction with a spectacle lens is usually impossible, but a point focus can be achieved with a rigid contact lens. Irregular astigmatism can be indicative of higher-order aberrations such as keratoconus or corneal ectasia.

The magnitude of corneal astigmatism has been reported to increase with age[6,7] and there is a shift from the steepest corneal meridian from the vertical (with-the-rule) to the horizontal meridian (against-the-rule)[8–11]. Data on the prevalence and severity of corneal astigmatism is typically obtained from case series of patients undergoing cataract surgery[11–16], with limited data from population based cross-sectional studies or large cohort studies[17].

The UKBB study[18–20] recruited over 500,000 men and women aged 40 to 69 years between 2006-2010 from the general population. In 2009 the study protocol was updated to include measurement of ocular data including corneal keratometry, on a subset of these[20]. The aim of our analysis is to describe corneal astigmatism and derived variables in the UKBB population, to look for associations with other biometric variables, socio-demographic factors, and eye conditions.

## Methodology

### UKBB Participants

The UK Biobank participants have previously been described in detail[20]. In brief, all adults aged between 49 and 69 years old who were registered with the UK National Health Service and living within 25 miles of one of the 22 participating study sites were invited to participate. From a total of 9.2 million postal invitations, 503,325 participants were recruited between 2006 and 2010 (response rate of 5.5%) and after accounting for withdrawals; data on 502,642 participants were available for analysis. All those recruited completed detailed questionnaires on their lifestyle, socioeconomic status, environment and health, and had a number of physiological measures from urine, saliva and blood samples. Further information can be found on the UK Biobank online data showcase (http://biobank.ctsu.ox.ac.uk/crystal/label.cgi).

### Ethics

All UK Biobank participants gave written, informed consent. The UK Biobank study was conducted under approval from the NHS National Research Ethics Service (Ref. 11/NW/ 0382), and anonymised data were provided from UK Biobank under application reference 10536.

### Eye measurements

Six of the recruiting centres performed an ophthalmic assessment[4] that included LogMAR visual acuity, autorefraction and keratometry (Tomey RC 5000 auto-ref-keratometer Tomey Corp., Nagoya, Japan), intraocular pressure (IOP) (Goldmann-correlated and Corneal-compensated) and corneal biomechanics (both Ocular Response Analyzer, Reichert, Depew, NY, USA). In total 117,279 (23.3%) of those enrolled had an ophthalmic assessment.

The Tomey RC 5000 examination produced 38 autorefraction and keratometric measurements for each eye (category: http://biobank.ctsu.ox.ac.uk/crystal/label.cgi?id=100014). This category includes data on whether the measurement was made, and the test result for refractometry (sphere, cylinder, axis, pupil diameter) and keratometry (corneal refraction and astigmatism). Corneal astigmatism was defined as the 3mm strong meridian minus the 3mm weak meridian. The average of these two values was defined as the mean corneal power. Similarly to[4], we defined spherical equivalent as the spherical power plus half of the refractive cylindrical power.

The Reichert Ocular Response Analyser (Reichert Corp., Philadelphia, PA), which measures the biomechanical distortion of the cornea produced by a puff of air, yielded a further 9 types of measurements for each eye (category: http://biobank.ctsu.ox.ac.uk/crystal/label.cgi?id=100015). These included eye applanation, corneal hysteresis, corneal resistance factor, corneal-compensated intraocular pressure, and an IOP graph for each eye. Participants who had eye surgery within the previous 4 weeks or those with possible eye infections did not have IOP measured.

### Self-reported eye conditions

The UKBB touchscreen questionnaire allowed participants to report if they wore glasses or contact lenses, and if they had an eye disorders or eye diseases, any injury or trauma, which eye was affected and when it was diagnosed. Refractive eye conditions included astigmatism, myopia, hyperopia, presbyopia, strabismus and amblyopia. Eye diseases include diabetic retinopathy, glaucoma, cataract or age-related macular degeneration.

### Socio-economic status and ethnicity

The Townsend deprivation index was determined using the participant’s postcodes at recruitment. The Townsend deprivation index has a UK mean of zero, with negative being less deprived and positive being more deprived. Ethnicity choices included white (English/Irish or other white), Asian or British Asian (Indian/Pakistani/Bangladeshi or other Asian), black or black British (Caribbean, African, or other black), Chinese, mixed (white and black Caribbean or African, white and Asian, or other mixed ethnicity), or other ethnic group (not defined).

### Lifestyle and environment

The UKBB touchscreen questionnaire also offered questions to participants about their lifestyle, health and environment. In particular, smoking status, alcohol intake frequency, use of sun/UV protection, skin colour without tanning, presence of diabetes. The possible questions/answers and their encoding are explained in more detail in the Table S1.

### Physical measures and disease

Blood pressure and heart rate were measured using the HEM-70151T digital blood pressure monitor (Omron, Hoofddorp, The Netherlands). Weight was measured with the BV-418 MA body composition analyzer (Tanita, Arlington Heights, IL). Height was measured using a Seca 202 height measure (Seca, Birmingham, UK). Body mass index (BMI) was calculated as weight in kg divided by height in m^2^. Waist circumference at the level of the umbilicus was measured using a Wessex non-stretchable tape measure. The UKBB includes diagnoses extracted from records across all their episodes in hospital coded using the International Classification of Diseases version-10 (ICD10).

### Participant selection

Of the 502642 participants in UK Biobank, 109,935 had 3mm strong or weak corneal meridian measurement values available for both eyes from which corneal astigmatism measurements could be derived. Participants were excluded if they had any of the following: previous laser refractive eye surgery (n=7440), previous eye surgery (for cataract, glaucoma or corneal graft) (n=8051), unreliable 3mm asymmetry index (n=12,910) or an unreliable keratometry result (n=6916). This left a total of 83,751 individuals for further analysis.

### Mean corneal power and axis of astigmatism

We tested the association of mean corneal power and axis of astigmatism with age and gender for both eyes. The axis of astigmatism was defined as the angle of the strong meridian minus the angle of the weak meridian. The axis of astigmatism was either 90 or −90 degrees for all individuals; this implies that no irregular astigmatism was detected among our participants.

### Corneal astigmatism, irregular corneal astigmatism and keratoconus

We explored the distribution of astigmatism and compared this to previously published studies. We then looked for patients with features suggestive of keratoconus such as mean corneal power exceeding 48, 49 or 50 dioptres[21,22]. The Tomey RC 5000 auto-ref-keratometer also provided an index irregular corneal astigmatism (normal, doubtful, high possibility of abnormality) and we therefore further refined our search for keratoconus using a steep corneal meridian of 48D or more, in the presence a “doubtful” or “high possibility of abnormality” 3mm index of irregular astigmatism.

### Statistical associations with corneal astigmatism, mean corneal power

Univariable linear regression and multivariable regression analysis models were applied to investigate predictors of corneal astigmatism. Variables were re-coded according to Table S1. To account for multiple testing, a Bonferroni corrected P value threshold of < 0.001 was applied to avoid false-positives due to the large number of tests carried out. Only parameters that showed significant association in the univariable analysis were included in the multivariable analysis. Since we found that corneal astigmatism measurements were asymmetric, with left eye having on average higher corneal astigmatism than right eye (Figure S1.a), we repeated statistical analysis in both eyes and only reported parameters which were consistently significantly associated in both eyes. We also repeated the statistical associations with a log-scaled corneal astigmatism variable since the p-value calculation in a linear regression assumes a normally distributed response variable (Figure S2.a.b). All analyses were performed using R statistical software version 3.2.3. The code is available at https://github.com/pontikos/UKBB/.

## Results

### Participant selection and distribution of corneal astigmatism

Of the 502,642 UKBB participants who had keratometry measures, after exclusions, 83,751 participants were selected for the purpose of this study. Of these, 36,490 (44%) were male. Ethnicity was 90% white, 3.44% Asian, 3.01% black, 0.89% mixed and 0.41% Chinese (Table 1). In the right eye, 69%, 46%, 29%, 11% and 5% had corneal astigmatism greater than or equal to 0.5, 0.75, 1.0, 1.5 and 2.0 dioptres respectively, and in the left eye, 69%, 46%, 30%, 11% and 5% (Figure S2.c). Anisometropia of less than 1 dioptre was found in 95% of individuals and a difference of more than 2 dioptres was found in the 0.83% of eyes. After stratification of participants by age group (decade) and gender, corneal astigmatism was found to decrease with age and to be on average higher in females than in males across age groups (Table 2).

**Table 1:** Distribution of participants in the UKBB across the different variables. Mean/sd or percentage of the 83,751 study participants in the UKBB by sex.

| variable | total | males | females | P value |
| --- | --- | --- | --- | --- |
| Age, years | 57.1 (8.1) | 57.3 (8.2) | 56.9 (8.0) | <.001 |
| Ethnicity white | 92.1 | 92.2 | 92 | <.001 |
| Ethnicity asian | 3.5 | 4 | 3.1 |  |
| Ethnicity black | 3.1 | 2.8 | 3.3 |  |
| Ethnicity mixed | 0.9 | 0.7 | 1 |  |
| Ethnicity chinese | 0.4 | 0.3 | 0.5 |  |
| Age completed full time education | 16.9 (2.5) | 16.9 (2.7) | 16.8 (2.3) | <.001 |
| Skin colour * | 2.3 (0.9) | 2.4 (0.9) | 2.3 (0.9) | <.001 |
| Use of UV protection | 2.7 (0.9) | 2.5 (0.9) | 2.9 (0.9) | <.001 |
| Alcohol intake ** | 3.0 (1.6) | 3.3 (1.5) | 2.8 (1.6) | <.001 |
| Season of assessment spring | 34.6 | 34.7 | 34.6 |  |
| Season of assessment autumn | 23.7 | 23.5 | 23.8 |  |
| Season of assessment winter | 22.1 | 22.4 | 21.9 |  |
| Season of assessment summer | 19.6 | 19.5 | 19.7 |  |
| Corneal corrected IOP, mmHg | 16.1 (4.2) | 16.4 (4.2) | 15.9 (4.3) | <.001 |
| Corrected Visual acuity, logMAR | 0.0 (0.2) | 0.0 (0.2) | 0.0 (0.2) | <.001 |
| Corneal resistance factor | 10.7 (2.3) | 10.6 (2.3) | 10.9 (2.4) | <.001 |
| Corneal hysteresis | 10.6 (2.3) | 10.4 (2.2) | 10.8 (2.3) | <.001 |
| Height, m | 168.3 (9.2) | 175.7 (6.8) | 162.7 (6.3) | <.001 |
| Weight, 10 kg | 77.4 (15.8) | 85.5 (14.2) | 71.3 (14.0) | <.001 |
| BMI, kg/m2 | 27.3 (4.8) | 27.7 (4.2) | 27.0 (5.1) | <.001 |
| SBP, mmHg | 139.7 (19.5) | 142.3 (18.3) | 137.6 (20.2) | <.001 |
| DBP, mmHg | 81.9 (10.6) | 83.6 (10.4) | 80.5 (10.5) | <.001 |
| Townsend deprivation index | -1.0 (3.0) | -1.0 (3.0) | -1.0 (2.9) |  |
| Smoker: true | 90.1 | 87.9 | 91.8 | <.001 |
| Smoker: false | 9.9 | 12.1 | 8.2 |  |
| Age Asthma Diagnosed, self reported | 30.5 (18.5) | 27.3 (19.1) | 32.7 (17.7) | <.001 |
| Diabetes, doctor diagnosed: true | 94.8 | 93.1 | 96.1 | <.001 |
| Diabetes, doctor diagnosed: false | 5.2 | 6.9 | 3.9 |  |

D = dioptres; \* Skin colour is coded as: very fair=1, fair=2, light olive=3, dark olive=4, brown=5, black=6;
\*\* Alcohol intake is coded as: never=0, special occasions=1, one to three times a month=2, once twice a week=3, three four times a week=4, daily=5.

**Table 2:** Mean, standard deviation, 25th and 75th percentile of right and left corneal astigmatism by age and gender of the 83,751 study participants in the UK BioBank. Corneal astigmatism decreases slightly with age and is slightly higher in females than in males. Left corneal astigmatism tends to be consistently higher than right corneal astigmatism (as illustrated in Figure S1.a).

| cohorts | right eye 3mm corneal astigmatism | left eye 3mm corneal astigmatism |
| --- | --- | --- |
| Men 40-49 | 0.861 (0.683, 0.430-1.090) | 0.864 (0.674, 0.440-1.100) |
| Men 50-59 | 0.810 (0.640, 0.400-1.030) | 0.818 (0.646, 0.400-1.050) |
| Men 60-69 | 0.788 (0.619, 0.390-1.000) | 0.792 (0.627, 0.400-1.000) |
| Women 40-49 | 0.923 (0.625, 0.510-1.190) | 0.942 (0.638, 0.520-1.200) |
| Women 50-59 | 0.890 (0.611, 0.470-1.140) | 0.915 (0.645, 0.490-1.170) |
| Women 60-69 | 0.850 (0.619, 0.440-1.090) | 0.855 (0.619, 0.440-1.090) |
| Total 40-49 | 0.896 (0.652, 0.470-1.150) | 0.908 (0.655, 0.480-1.160) |
| Total 50-59 | 0.857 (0.624, 0.440-1.100) | 0.875 (0.647, 0.450-1.120) |
| Total 60-69 | 0.822 (0.620, 0.420-1.050) | 0.827 (0.623, 0.420-1.050) |
| All | 0.848 (0.627, 0.430-1.090) | 0.857 (0.636, 0.440-1.100) |
| Difference (left-right) | -0.009 (-0.026 0.01), P=0.340730 |  |

### Association of mean corneal power, axis of astigmatism and anisometropia with age and gender

Older age was significantly associated with increased mean corneal power in both eyes, with an average increase of 0.015 (0.014 to 0.016) dioptres per year (Figure S3.b). Axis of astigmatism changed with older age from with-the-rule (90 degrees) to against-the-rule (−90 degrees) (Figure S3.c). Anisometropia was slightly more prevalent in males than in females (0.98% vs 0.71%).

### Irregular corneal astigmatism and keratoconus

The number of individuals with a strong 3mm corneal meridian greater than 48, 49 and 50 dioptres, in the presence of “doubtful” or “high possibility of abnormality” index of irregular corneal astigmatism in both eyes, were 132, 46 and 5 respectively (less than 0.16% of the cohort). However, none of these individuals had a keratoconus diagnosis (H18.6) in the UKBB. Of the 502,642 in the UKBB, only 9 had a keratoconus diagnosis, none of which were in our filtered list of 83,751 participants. Four of these individuals were filtered out because they previously had surgery, the other five were not included because they had no keratometry values.

### Univariable and multivariable analysis of corneal astigmatism

By order of magnitude, the univariable analysis found that - decreased corrected visual acuity, Asian and black ethnicity, male gender, darker skin colour, decreased use of UV protection, increased alcohol intake, increased corneal corrected IOP, older age and younger age completed full-time education – were significantly associated with decreased corneal astigmatism (Table 3.1 and Table S2.1). After including these variables in the multivariable analysis (Table 3.2 and Table S2.2), the following parameters remained significantly associated with decreased corneal astigmatism: decreased corrected visual acuity, male gender, darker skin colour, increased alcohol intake, older age and younger age completed full-time education.

**Table 3.1.**
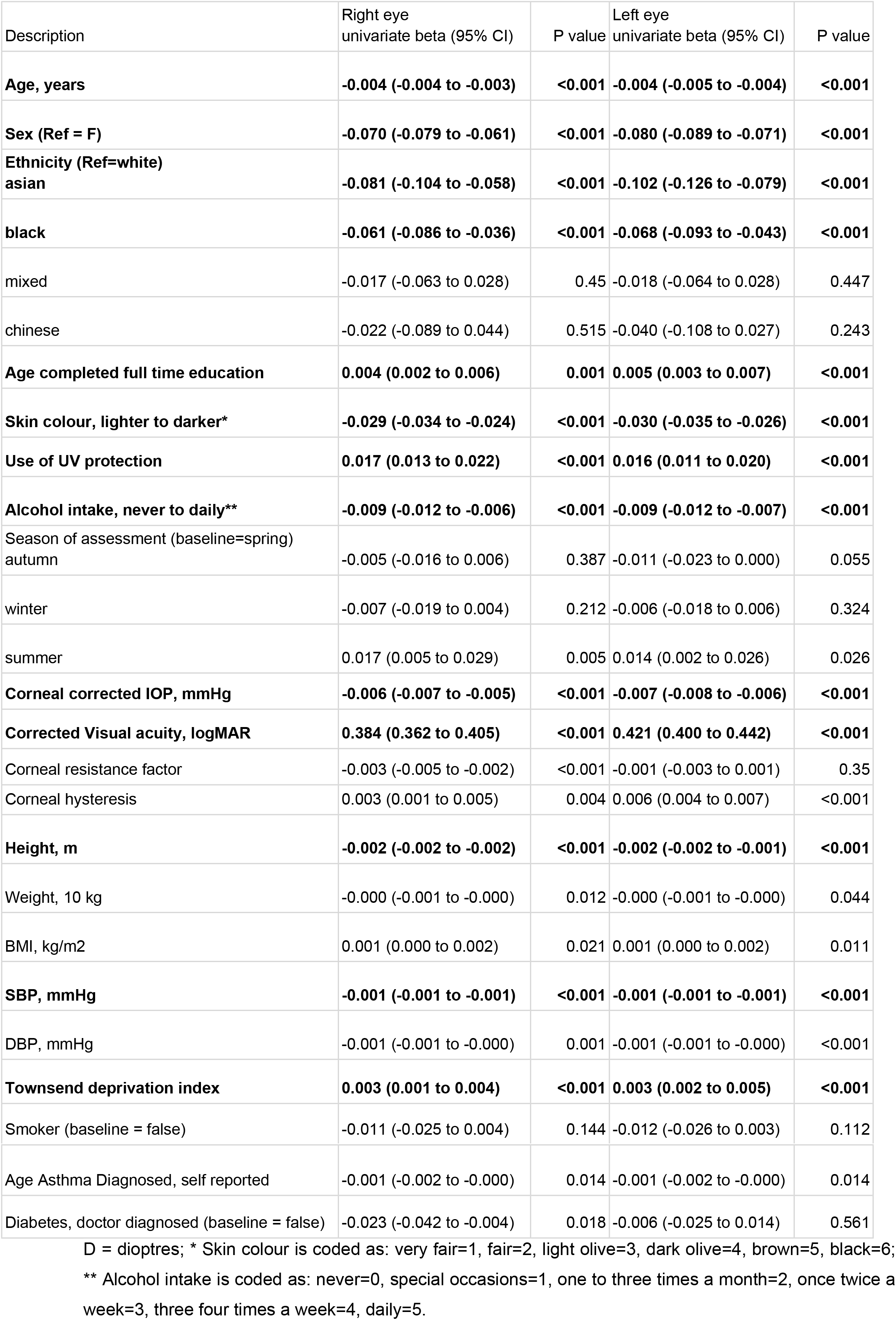
Results of univariable regression in 83,751 study participant in the UKBB for right and left eye corneal astigmatism. Significant associations are highlighted in bold.

| Description | Right eye<br>univariate beta (95% CI) | P value | Left eye<br>univariate beta (95% CI) | P value |
| --- | --- | --- | --- | --- |
| **Age, years** | **-0.004 (-0.004 to -0.003)** | **<0.001** | **-0.004 (-0.005 to -0.004)** | **<0.001** |
| **Sex (Ref = F)** | **-0.070 (-0.079 to -0.061)** | **<0.001** | **-0.080 (-0.089 to -0.071)** | **<0.001** |
| **Ethnicity (Ref=white)**<br>**asian** | **-0.081 (-0.104 to -0.058)** | **<0.001** | **-0.102 (-0.126 to -0.079)** | **<0.001** |
| **black** | **-0.061 (-0.086 to -0.036)** | **<0.001** | **-0.068 (-0.093 to -0.043)** | **<0.001** |
| mixed | -0.017 (-0.063 to 0.028) | 0.45 | -0.018 (-0.064 to 0.028) | 0.447 |
| chinese | -0.022 (-0.089 to 0.044) | 0.515 | -0.040 (-0.108 to 0.027) | 0.243 |
| **Age completed full time education** | **0.004 (0.002 to 0.006)** | **0.001** | **0.005 (0.003 to 0.007)** | **<0.001** |
| **Skin colour, lighter to darker\*** | **-0.029 (-0.034 to -0.024)** | **<0.001** | **-0.030 (-0.035 to -0.026)** | **<0.001** |
| **Use of UV protection** | **0.017 (0.013 to 0.022)** | **<0.001** | **0.016 (0.011 to 0.020)** | **<0.001** |
| **Alcohol intake, never to daily\*\*** | **-0.009 (-0.012 to -0.006)** | **<0.001** | **-0.009 (-0.012 to -0.007)** | **<0.001** |
| Season of assessment (baseline=spring)<br>autumn | -0.005 (-0.016 to 0.006) | 0.387 | -0.011 (-0.023 to 0.000) | 0.055 |
| winter | -0.007 (-0.019 to 0.004) | 0.212 | -0.006 (-0.018 to 0.006) | 0.324 |
| summer | 0.017 (0.005 to 0.029) | 0.005 | 0.014 (0.002 to 0.026) | 0.026 |
| **Corneal corrected IOP, mmHg** | **-0.006 (-0.007 to -0.005)** | **<0.001** | **-0.007 (-0.008 to -0.006)** | **<0.001** |
| **Corrected Visual acuity, logMAR** | **0.384 (0.362 to 0.405)** | **<0.001** | **0.421 (0.400 to 0.442)** | **<0.001** |
| Corneal resistance factor | -0.003 (-0.005 to -0.002) | <0.001 | -0.001 (-0.003 to 0.001) | 0.35 |
| Corneal hysteresis | 0.003 (0.001 to 0.005) | 0.004 | 0.006 (0.004 to 0.007) | <0.001 |
| **Height, m** | **-0.002 (-0.002 to -0.002)** | **<0.001** | **-0.002 (-0.002 to -0.001)** | **<0.001** |
| Weight, 10 kg | -0.000 (-0.001 to -0.000) | 0.012 | -0.000 (-0.001 to -0.000) | 0.044 |
| BMI, kg/m2 | 0.001 (0.000 to 0.002) | 0.021 | 0.001 (0.000 to 0.002) | 0.011 |
| **SBP, mmHg** | **-0.001 (-0.001 to -0.001)** | **<0.001** | **-0.001 (-0.001 to -0.001)** | **<0.001** |
| DBP, mmHg | -0.001 (-0.001 to -0.000) | 0.001 | -0.001 (-0.001 to -0.000) | <0.001 |
| **Townsend deprivation index** | **0.003 (0.001 to 0.004)** | **<0.001** | **0.003 (0.002 to 0.005)** | **<0.001** |
| Smoker (baseline = false) | -0.011 (-0.025 to 0.004) | 0.144 | -0.012 (-0.026 to 0.003) | 0.112 |
| Age Asthma Diagnosed, self reported | -0.001 (-0.002 to -0.000) | 0.014 | -0.001 (-0.002 to -0.000) | 0.014 |
| Diabetes, doctor diagnosed (baseline = false) | -0.023 (-0.042 to -0.004) | 0.018 | -0.006 (-0.025 to 0.014) | 0.561 |

D = dioptries; \* Skin colour is coded as: very fair=1, fair=2, light olive=3, dark olive=4, brown=5, black=6; \*\* Alcohol intake is coded as: never=0, special occasions=1, one to three times a month=2, once twice a week=3, three four times a week=4, daily=5.

**Table 3.2.** Results of multivariable regression in 83751 study participant in the UKBB for right and left eye corneal astigmatism. Only parameters that were significant in the univariable regression were included in the multivariable regression. Significant associations are highlighted in bold.

| Description | Right eye<br>multivariate beta (95%<br>CI) | P value | Left eye<br>multivariate beta (95% CI) | P value |
| --- | --- | --- | --- | --- |
| **Age, years** | **-0.006 (-0.006 to -0.005)** | **<0.001** | **-0.006 (-0.006 to -0.005)** | **<0.001** |
| **Sex (Ref = F)** | **-0.058 (-0.075 to -0.042)** | **<0.001** | **-0.082 (-0.099 to -0.066)** | **<0.001** |
| Ethnicity (Ref=white)<br>asian | -0.043 (-0.083 to -0.003) | 0.034 | -0.083 (-0.124 to -0.043) | <0.001 |
| black | 0.003 (-0.043 to 0.049) | 0.896 | -0.033 (-0.079 to 0.014) | 0.165 |
| mixed | -0.071 (-0.133 to -0.009) | 0.025 | -0.030 (-0.092 to 0.033) | 0.356 |
| chinese | -0.100 (-0.201 to 0.002) | 0.054 | -0.085 (-0.186 to 0.017) | 0.102 |
| **Age completed full time education** | **0.006 (0.004 to 0.008)** | **<0.001** | **0.007 (0.005 to 0.009)** | **<0.001** |
| **Skin colour, lighter to darker\*** | **-0.032 (-0.041 to -0.023)** | **<0.001** | **-0.027 (-0.036 to -0.018)** | **<0.001** |
| Use of UV protection | 0.010 (0.003 to 0.016) | 0.003 | 0.003 (-0.004 to 0.009) | 0.379 |
| **Alcohol intake, never to daily\*\*** | **-0.007 (-0.010 to -0.003)** | **<0.001** | **-0.007 (-0.011 to -0.003)** | **<0.001** |
| Corneal corrected IOP, mmHg | -0.004 (-0.007 to -0.001) | 0.009 | **-0.006 (-0.007 to -0.004)** | **<0.001** |
| **Corrected Visual acuity, logMAR** | **0.441 (0.413 to 0.469)** | **<0.001** | **0.476 (0.449 to 0.504)** | **<0.001** |
| Corneal resistance factor | -0.009 (-0.019 to 0.002) | 0.096 |  |  |
| Corneal hysteresis | 0.003 (-0.009 to 0.015) | 0.621 | -0.003 (-0.006 to -0.001) | 0.019 |
| Height, m | -0.000 (-0.001 to 0.001) | 0.65 | 0.001 (-0.000 to 0.001) | 0.203 |
| SBP, mmHg | -0.000 (-0.000 to 0.000) | 0.773 | -0.000 (-0.001 to 0.000) | 0.12 |
| DBP, mmHg | 0.001 (0.000 to 0.002) | 0.022 | 0.001 (0.000 to 0.002) | 0.024 |
| Townsend deprivation index | 0.002 (0.000 to 0.004) | 0.018 | 0.002 (0.000 to 0.004) | 0.02 |

D = dioptres; \* Skin colour is coded as: very fair=1, fair=2, light olive=3, dark olive=4, brown=5, black=6;
\*\* Alcohol intake is coded as: never=0, special occasions=1, one to three times a month=2, once twice a week=3, three four times a week=4, daily=5.

## Discussion

### Distribution of corneal astigmatism in the UKBB compared to other cohorts

The distribution of astigmatism in the large population reported in this study supports evidence from previous smaller studies, both in the UK and worldwide, in pre-operative patients [X1] and from large consortiums such as CREAM (n=55,177)[17]. We found that 69%, 29%, 11% and 5% had corneal astigmatism ≥0.5, 1.0, 1.5 and 2.0 dioptres respectively. These are slightly lower than values reported from a recent series of 110,468 cataract pre-operative eyes [23] where 78%, 42%, 21% and 11% having corneal astigmatism ≥0.5, 1.0, 1.5 and 2.0 dioptres respectively. A study of 1,230 eyes undergoing cataract surgery in Wales found corneal astigmatism of >0.5D in 75% in Wales[15] (N=1,230 eyes). Corneal astigmatism ≥1.0D was found in 36% of eyes with cataract in Germany[24] (N=15,448 eyes), 47% in China (N=12,449)[25] and 35% in South Korea[26] (N=2,847 eyes). Recently, Curragh et al[27] reported that 41% of eyes undergoing cataract surgery (N=2,080) in Northern Ireland had >1.0D of corneal astigmatism. However cataract surgery is usually performed in an older age group than those of the participants in the UK Biobank and these pre-operative clinical groups are not necessarily comparable to UKBB participants. A recent CREAM study[28] reported the median corneal astigmatism and median age across 22 studies (8 Asian and 14 European). The median corneal astigmatism was reported in each study and this ranged from 0.539D in the Rotterdam-II European study (N=3964, mean age=64.8), to 1.21D in the Asian Singapore Cohort Study of the Risk Factors for Myopia (SCORM) study (N=1894, mean age=10.8). Comparable studies to the UKBB in terms of age and gender demographics of the participants are the Rotterdam-III Study (N=5850 eyes, mean age=57)[29], the Singapore Chinese Eye Study (SCES-610K) (N=1106 eyes, mean age=57.6)[30], the Gutenberg Health Study (GHS-1) study (N=4796 eyes, mean age=55.9)[31] which reported a median corneal astigmatism of 0.618D, 0.703D and 0.65D respectively. This is comparable to the UKBB median corneal astigmatism of 0.71D.

### Modelling of corneal astigmatism

In the multivariable analysis, parameters known to be strongly associated with gender, such as height and weight (Table 3.2), were no longer significantly associated with corneal astigmatism. The adjusted R-squared of the multivariable regression was remarkably low at 0.015 which highlights that there are many other unobserved variables which influence corneal astigmatism.

### Corneal astigmatism, amblyopia and strabismus

The number of eyes affected by amblyopia and strabismus in the UKBB are 2483 and 1052 respectively. Corneal astigmatism is highest in eyes affected by amblyopia and strabismus (Figure S4.a). This confirms that, as previously reported by [32,33], uncorrected high corneal astigmatism is a significant risk factor for amblyopia (OR=1.98 (1.87 to 2.09), P<0.001) and strabismus (OR=1.73 (1.59 to 1.88), P<0.001) (Figure S4.b).

### Corneal astigmatism and IOP measurements

Of interest, corneal hysteresis, which measures the cornea’s ability to absorb and dissipate energy, was not found to be associated with corneal astigmatism in the univariable or multivariable analysis. However when corneal astigmatism was log-transformed, a small but positive significant association was detected for both eyes in the univariable analysis (Table S2.1) but not in the multivariable analysis (Table S2.2). These results suggest that the ability of the cornea to absorb and dissipate energy (corneal hysteresis) was not strongly significantly associated with corneal astigmatism. However, corneal hysteresis is significantly positively associated with mean corneal power in both eyes. In the univariable analysis (Table 3.1), we found a small by significant negative association between corneal corrected intraocular pressure and corneal astigmatism in both eyes, however in the multivariable analysis the association was no longer significant in both eyes (Table 3.2).

### Corneal astigmatism and gender

Our study confirms as previously reported by[25], that corneal astigmatism is higher in females than in males even after adjusting for weight and height (Table 3.2). Males have on average 0.06D less corneal astigmatism in right eye than females and 0.08D less in left eye.

### Corneal astigmatism and age

Corneal astigmatism decreases significantly with age by an average 0.06D in both eyes per decade even after controlling for weight and height (Table 3.2). This is contrary to what is reported by[34] namely that the level of corneal astigmatism is relatively constant across age. As reported by Shah et al in a previous UKBB analysis[34], we did confirm the stronger association between age and refractive (cylindrical) astigmatism in right eye (B=0.011 (0.010 to 0.011), P<.001) and left eye (B=0.010 (0.09 to 0.010), P<0.001) (Figure S3.a). The fact that refractive astigmatism increases with age but that corneal astigmatism decreases with age suggest that lenticular astigmatism may be driving the increase in refractive astigmatism, possibly due to the onset of presbyopia. We also found that age is significantly positively associated with corneal power both in right (B=0.015 (0.014 to 0.016), P<0.001) and left eye (B=0.015 (0.014 to 0.016), P<0.001) (Figure S3.b). We also observed that the axis of astigmatism changes with age from with to against the rule (Figure S3.c) significantly in the right eye but not in the left eye.

### Corneal astigmatism and age completed full-time education

We discovered a significant positive association between age at which full-time education was completed and corneal astigmatism (B=0.006 (0.004 to 0.008)). This result was consistent with participants with self-reported astigmatism finishing full-time education later than other participants (Figure S4.c). Interestingly, this relationship was not observed in individuals with myopia (Figure S4.c). In fact, recent evidence suggests that myopia is perhaps not linked as much to poor lighting conditions and near work[35], but rather to earlier life exposures [36].

### Corneal astigmatism, ethnicity and skin colour

Asian and black ethnicities appear to be significantly protective for corneal astigmatism in both eyes according to the univariable analysis (Table 3.1 and S3.1), but are no longer significant in the multivariable analysis (Table 3.2 and S3.2) for log transformed corneal astigmatism. However skin colour remains significantly associated with darker skin being protective (B=−0.032) (Table 3.2 and S3.2). This relationship can also be clearly seen in Figure S5. The link between corneal astigmatism and deficiency in melanin production has been previously reported for albinism[37] but we have now also reported this association in a healthy population via darkness of skin showing that darker skin and hence increased melanin production appears protective for corneal astigmatism.

### Corneal astigmatism and alcohol intake

Alcohol intake is significantly protective for corneal astigmatism according to the univariable and multivariable analysis. In particular, the group that drink nearly every day has the lowest corneal astigmatism. This is surprising due to the negative consequences of alcohol abuse on eye conditions. However on closer inspection it appears that the group that drinks nearly every day in the UKBB consists primarily of men in the 65+ age group; 55% of men drink every day in this study vs 44% of women. It then follows that alcohol intake effect is difficult to decouple from gender due to the high correlation. In fact, the three-way interaction between alcohol-intake, age and gender is illustrated in Figure S6, with “never-drinkers” and “daily drinkers” showing a clear interaction.

### Corneal astigmatism and keratoconus

We looked for participants with features suggestive of keratoconus such as mean corneal power exceeding 48, 49 or 50 dioptres[21,22] and also explored the Tomey RC 5000 auto-ref-keratometer specific parameter of irregular corneal astigmatism as a proxy for a keratoconus diagnosis. However none of the 83,751 participants had a prior diagnosis of keratoconus (H18.6). In the entire UKBB, only 9 participants had a keratoconus diagnosis, none of these were in our included list of 83,751 participants due to previous exclusion as detailed earlier. This number (1:10,000) is a factor of 10 less than population-based estimates from North Europe [38]. Based on these findings, identification of keratoconus in cross-sectional or population studies would appear to require corneal topography or corneal tomography map review rather than evaluation of simple keratometry indices.

### Strengths and limitations of our study

The strength of this study is the large sample size of 83,751 participants and that participants were not pre-operative patients hence more representative of the general population. However due to the limited age range of the participants, between 40 and 69 years, and the voluntary nature of study participation, the age distribution is not representative of the UK and participants are likely to be a healthier more educated sample of the UK population. Regardless, a range of exposures and characteristics are likely to have been captured due to the sample size and so the results can still be applicable to other populations.

## Conclusion

In conclusion, this analysis of associations with astigmatism in a large cohort confirms previous associations including age and gender, and identified novel associations including age completed full time education, skin colour and alcohol intake. The highest risk group for corneal astigmatism were younger females of lighter skin colour, having completed full time education later, with higher corrected visual acuity. It was also confirmed that uncorrected corneal astigmatism is a high-risk factor for amblyopia and strabismus.

## Acknowledgments

This research has been conducted using the UK Biobank Resource under Application Number 10536. Collaborators on the application are Nikolas Pontikos, Alexander Day, Parul Desai, Paul Foster and Stephen Tuft. The PI is Alexander Day. The collection of eye & vision data in UK Biobank was supported in part by a grant from the NIHR Biomedical Research Centre at Moorfields Eye Hospital and UCL Institute of Ophthalmology. The UK Biobank Eye and Vision Consortium is supported by a grant from the Special Trustees of Moorfields Eye Hospital (now Moorfields Eye Charity). The main contact for this consortium is Prof Paul Foster and list of members is available from the consortium website (http://www.ukbiobankeyeconsortium.org.uk/people).

## Author Contributions

Conceived and designed the experiments: ACD, NP and SJT. Analysed the data: NP. Wrote the paper: NP, ACD, SJT and PJF. Review and critique of the manuscript: ACD, SC, PJF and SJT.

